# PhyloPi: an affordable, purpose built phylogenetic pipeline for the HIV drug resistance testing facility

**DOI:** 10.1101/367946

**Authors:** Phillip A Bester, Andrie De Vries, SJPK Riekert, Kim Steegen, Gert van Zyl, Dominique Goedhals

## Abstract

Phylogenetic analysis plays a crucial role in quality control in the HIV drug resistance testing laboratory. If previous patient sequence data is available sample swaps can be detected and investigated. As Antiretroviral treatment coverage is increasing in many developing countries, so is the need for HIV drug resistance testing. In countries with multiple languages, transcription errors are easily made with patient identifiers. Here a self-contained blastn integrated phylogenetic pipeline can be especially useful. Even though our pipeline can run on any unix based system, a Raspberry Pi 3 is used here as a very affordable and integrated solution.

The computational capability of this single board computer is demonstrated as well as the utility thereof in the HIV drug resistance laboratory. Benchmarking analysis against a large public database shows excellent time performance with minimal user intervention. This pipeline also contains utilities to find previous sequences as well as phylogenetic analysis and a graphical sequence mapping utility against the *pol* area of the HIV HXB2 reference genome. Sequence data from the Los Alamos HIV database was analyzed for inter- and intra-patient diversity and logistic regression was conducted on the calculated genetic distances. These findings show that allowable clustering and genetic distance between viral sequences from different patients is very dependent on subtype as well as the area of the viral genome being analyzed.

The Raspberry Pi image for PhyloPi, source code of the pipeline, sequence data, bash-, python- and R-scripts for the logistic regression, benchmarking as well as helper scripts are available at http://scholar.ufs.ac.za:8080/xmlui/handle/11660/7638 and https://github.com/ArmandBester/phylopi.

The PhyloPi image and the source code are published under the GPLv3 license. A demo version of the PhyloPi pipeline is available at http://phylopi.hpc.ufs.ac.za/.

## Introduction

The use of combined Antiretroviral Therapy (cART) has dramatically decreased the mortality and morbidity of HIV infected people. It was estimated that 12.9 million people were receiving antiretroviral treatment (ART) by the end of 2013 worldwide. The number of HIV infected individuals not receiving ART dropped from 90% in 2006 to 63% in 2013 globally [1]. Since 2010 the number of people living with HIV on antiretroviral therapy increased from 7.5 million to 17 million in 2015, whereas people infected with HIV only increased from 33.3 million to 36.7 million in the same time period [2]. However, inadequate adherence on ART, especially when low genetic barrier regimens such as NNRTI-based combination therapies are prescribed, often results in virologic failure with the emergence of drug resistance. In contrast, in cases with virologic failure of high genetic barrier regimens, such as boosted protease inhibitor- or dolutegravir-containing regimens drug resistance is often absent [2,3]. Drug resistance testing can differentiate patients, with persisting failure, who require adherence support from those who require a regimen switch and to select appropriate third-line regimens [4,5]. It has been demonstrated by numerous investigators that not only is drug resistance on the rise in South Africa and other countries, but that transmitted resistance is also on the increase (Manasa *et al.*, 2016; Huang *et al.*, 2009; Rhee *et al.*, 2015; Boender *et al.*, 2015; Steegen *et al.*, 2016a; Steegen *et al.*, 2016b). As a result, this will increase the amount of sequence data which will become available. We believe the work done here can help individual drug resistance facilities to cope with the quality assurance requirements this increase will infer.

Phylogenetic analysis is typically used in molecular HIV drug resistance testing facilities to detect gross cross contamination during nucleic acid extraction, reverse transcription, polymerase chain reaction or sequencing. In some cases, samples swaps can be detected if the patient has more than one viral sequence. However, in busy clinics and diagnostic facilities transcription errors can be made and errors are introduced in patient identification information, like names and date of birth. Also, from experience in our South African setting, patients can have multiple names which can be used interchangeably based on the ethnicity and language preference of the healthcare worker/patient pair. This makes it difficult and laborious to find previous viral sequences to use in phylogenetic analysis. Our pipeline includes a self-updating blast database which is used to find most similar past sequences to include in the phylogenic analysis.

Although, the comparison of current and past viral sequences does not guarantee that no sample swaps occurred, it does offer a valuable screen. The ability to include previous isolates in the drug resistance genotypic inference provides the resistance mutation history of the patient. It is probable that after treatment changes the circulating viral population changed and previous resistance mutations have reverted or have been displaced by wild type amino acid [11,12]. The archived proviral resistance can become important when formulating a 3^rd^ line treatment regimen in most third world settings.

Phylogenetic inference can be broken down into 4 consecutive steps [13]. The first step is a multiple alignment of sequences. This alignment is then curated, especially to look for and exclude alignment overhangs and in some cases gaps. This can dramatically improve the accuracy of the phylogenetic inference [14]. The curated alignment is then used as the input data for phylogenetic analysis and the phylogenetic tree is finally rendered. Combining these steps into an automated pipeline has opened phylogenetic analysis to the non-specialist and has created a convenient solution even for the specialist. This is very well demonstrated by the popular Phylogeny.fr web server, available at http://www.phylogeny.fr/index.cgi [13]. Also, the plethora of software available to conduct phylogenetic analysis can be daunting to the casual user as can be appreciated from Felsentein’s inventory available at http://evolution.genetics.washington.edu/phylip/software.html. Another very well cited and freely available software package for phylogenetic analysis is Mega [15]. The one thing which these ‘Swiss army knife’ implementations have in common is that already well-established and -cited software or algorithms are used to conduct the various steps. We demonstrate a purpose build, ultra-low cost, self-contained and portable solution to phylogenetic inference in the HIV drug resistance genotyping laboratory. Our pipeline is implemented on a Raspberry Pi 3. These pocket sized single board computers are the brain child of Eben Upton and colleagues form the Cambridge University Computer Laboratory [16]. We furthermore conducted logistic regression on publically available HIV sequence data from the Los Alamos HIV sequence database in order to demonstrate the variations between inter- and intra-patient genetic distances.

## Usage and general description

When powered on, PhyloPi broadcasts a WiFi hotspot to which another device can connect. It serves a simple web interface which is used as a graphical user front end. Here the user can set up the initial BLAST [17,18] database by formating a blastn database from a cumulative fasta file of all viral sequences sequencing data done previous. This requires the user to click on the “BLASTdb conf” button which then takes the user to the blast database setup page. Here the user uses the “Browse…” button to select the fasta file to be formatted into a blastn database. The user supplies a name for the database and click on “Format DB”. After this initial step the BLAST database gets updated every time the user submits new sequences for analysis, thus maintaining a rolling, up to date database. Before uploading the fasta file for phylogenetic inference, the user can choose how many most similar sequences from the BLAST sequence database must be included for analysis. The result for every analysis is stored in a SQL database which makes it possible to retrieve previous analysis as a compressed archive.

These past results can either be queried by a query sample used in the analysis, the subject sample found by blastn in any specific analysis or by name of the analysis. If all fields are left empty, all previous analyses will be listed. Every step of the pipeline is stored in a compressed result folder which allows the user to intervene on the phylogenetic inference from any step and to inspect each step. The phylogenetic tree rendering is available in portable network graphics (PNG) format, scalable vector graphics (SVG) format and portable document format (PDF). Also included in the result folder is a HTML R rendered heatmap showing the isolates in a distance matrix. This distance matrix is also available as a comma separated values file. Other files in the results folder include a fasta file containing the query sequences submitted for analysis combined with the subject samples retrieved by blastn, as well as the MAFFT [19] aligned fasta files pre and post TrimAl [14] curation. TrimAl also outputs a HTML file which annotates its actions on the MAFFT alignment; this is also included in the results archive.

The user can search for previous isolates by entering a comma separated list of full or partially matched identifiers depending and how they set a check box. This makes it easy to retrieve previous samples in case the user is interested in the resistance profile of the previous isolates.

Also, a ‘sanity check’ has been implemented which the user can use to determine if the input sequences are suitable for phylogenetic inference, or more generally are the sequences capable of generating a sensible alignment. It is conceivable, that in a busy drug resistance testing facility, the user might end up with sequences in a file which cover areas of the HIV genome which does not overlap. PhyloPi will not be able to handle such data and will fail after the TrimAl step. In such instance, PhyloPi warns the user about this and the offending sequence identifier is displayed in red. The ‘sanity check’ provides a crude way of mapping the individual entries in the fasta file to the POL region of HXB2 (Fig 1) which is available as one of the links on the results page. The ‘sanity check’ is also available as a separate mapping only tool through the link provided on the main page. Since blastn is already available in PhyloPi, it was used for the mapping against the HXB2 reference. From Fig 1, it is easy to find the sequences not suitable for inclusion in the phylogenetic inference by PhyloPi or any software. The sequence covering the protease only region cannot produce a sensible alignment with the sequences covering the reverse transcriptase only. A sequence covering the integrase area was also included for illustrative purposes. This graphic is rendered in SVG format, thus allowing very high resolution. This is important if the user has a fasta file with many entries. We have tested this on 105 sequences and were able to zoom in and quickly find sequences not making sense in the alignment.

**Fig 1.** A graphical representation of input sequences mapped against the reference, HXB2. The legend bar at the bottom shows the protease, reverse transcriptase and integrase in green, blue and red respectively. It is clear from this that the selected sequences do not qualify for phylogenetic inference.

### Considerations on software implemented

When choosing software for this pipeline the following factors were considered. The source code of the software must be available to allow for compilation. This is essential in order to obtain binary executables capable of running on the Raspberry Pi 3 ARM Cortex-A53 processor. The openness and availability of source code also provides transparency and the possibility of scrutiny, thus no ‘black boxes’. Since the Raspberry Pi 3 has only 1 GB of random access memory, the software used must have a lightweight memory footprint. For the computationally intensive tasks we selected software capable of parallelizing in order to maximize the use of the ARM Cortex-A53 quad core processor. Lastly, we only used well cited software with a command line interface.

## The pipeline

When the user uploads a new fasta file for analyses our pipeline first determines whether any unique entries were uploaded not yet present in the master fasta file. If a unique header is found in the fasta entry then that sequence is added to the master fasta file. Thus, if two or more sequences are identical, but the headers differ they will be added to the master fasta file. After new sequences have been added to the master fasta file, it gets formatted to a nucleic acid BLAST database with makeblastdb [18]. This updated BLAST database can now be used to retrieve the most relevant sequences for the analysis. In case different sequences as queries retrieve the same result sequence, these duplicates are removed. This combined input and blast retrieved sequences are saved as a fasta file which is used as the input for the multiple sequence alignment. The alignment is conducted using MAFFT [19], this is followed by removal of overhangs in the multiple sequence alignment using TrimAl [14]. The phylogenetic inference is conducted by FastTree 2 [20] and finally the ETE3 Python API (Application Program Interface) is used for tree rendering [21]. A block diagram of the PhyloPi pipeline is illustrated in Fig 2. The R programming language is used to calculate the genetic distances from the MAFFT aligned multiple sequence alignment using the packages seqinr and APE [22,23].

**Fig 2.** A block diagram illustrating the PhyloPi pipeline. A web interface offers control over the pipeline. With each sequence submission the master fasta file is updated and a BLAST database is formatted, thus maintaining a rolling blastn database. Duplicate entries are removed from the combined input and BLAST retrieved isolates and this is aligned with MAFFT. Overhangs are removed from the alignment before phylogenetic inference with FastTree. Finally, the tree is rendered with ETE3.

The heatmap is available in a comma separated values file as well as an interactive HTML file with colours corresponding to the genetic distances. The blocks of the heatmap are shaded from green to yellow to red and provide information such as sequence identifiers and genetic distances. The heatmap gives additional support to find close clustering of sequences. The heatmap colours are scaled such that closely related isolates are coloured red and distant related isolated are coloured green. This interactive HTML heatmap is rendered using the plotly R package.

As can be seen from Fig 2, the entire pipeline is completely self-contained and the only additional requirement is a WiFi capable device to be used for input. This provides extra security, in that the data is not available on the internet. Other noteworthy software used are shown in Table 1.

**Table 1.** Other software used for PhyloPi.

| Software | Use/Purpose | Link to software |
| --- | --- | --- |
| Hostapd and udhcpd | WiFi hotspot and DHCP server. Enables devices to connect to PhyloPi. | https://w1.fi/hostapd/<br>and<br>https://udhcp.busybox.net/ |
| Lighttpd 1.4.35 | Web server. | https://www.lighttpd.net/ |
| SQLite 2.8.17 | SQL database engine. To store results. | https://www.sqlite.org/ |
| R 3.3.3 | R | https://www.r-project.org/ |
| Open MPI | Message Passing Interface for mutli threading. | https://www.open-mpi.org/ |
| Python 2.7.9 | Used for the pipeline. | https://www.python.org/ |
| Xvfb and PyVirtualDisplay | X11 display server implementation for performing graphical operations and a python wrapper. | https://www.x.org/archive/X11R7.6/doc/man/man1/Xvfb.1.xhtml<br>and<br>https://pypi.python.org/pypi/PyVirtualDisplay |

## Application of PhyloPi in the HIV drug resistance testing laboratory

The HIV genotyping drug resistance facility at Department of Medical Microbiology and Virology, University of the Free State / National Health Laboratory Services has been using the PhyloPi pipeline for approximately a year. The following case reports are examples of applications of the PhyloPi in this facility where situations were resolved using this tool.

### Case 1

A sample for a HIV drug resistance test was processed and released for patient A (adult female) without any quality control problems. About a month and a half later a sample from the same clinic was processed for patient B. Our analysis with PhyloPi revealed a genetic distance of 0 between patient A and B. The requesting clinician was contacted and the situation explained. The clinician reported that patient A was seen at the clinic while patient B’s folder was still on the desk after review. When a nurse fetched patient A for phlebotomy the wrong file (that of patient B) was taken as the file for patient A, thus the identity of patient B was used for patient A on the sample. If only per-batch phylogenetic analysis were performed here, this mistake would have been missed.

### Case 2

A sample was processed for HIV drug resistance testing before the viral load which was taken on the same day was available. When the viral load became available it was <150 RNA copies/mL. The HIV drug resistance sample was resubmitted for viral load testing to exclude a sample swap during viral load testing; the same viral load result was however reported. Analysis with PhyloPi revealed a genetic distance of 0.006 in the genetic distance heatmap with a sample processed approximately 9 months earlier. Even though the name of the patient was spelled differently, the date of birth matched as well as the address. We thus concluded both samples belonged to the same patient.

### Vertical transmission and archived resistance

Also, mother and child pairs can be found with PhyloPi. Although this does not have obvious significant clinical meaning it does show that the pipeline works. In cases where patients have been genotyped before, it is possible that the patient’s viral population has lost drug resistance mutations due to the lack of selective pressure because of regimen changes. Here it can become important to include previous results, as it will give the clinician an idea of the archived drug resistance. Either way, PhyloPi reliably detects previously samples from same patients or mother to child transfer.

## Performance benchmarks

In order to demonstrate the capability of this small single board computer we performed benchmarks in terms of the time each step in the pipeline takes. Instead of using CPU time to illustrate the performance of the software used, we rather gathered the real time required to complete each step. Thereby we show the time requirements in relevance to the workings of the laboratory.

We downloaded HIV-1 sequences of the POL CDS region from the Los Alamos HIV public database (http://www.hiv.lanl.gov/). At the time of this analysis, 11 337 sequences were available. A descriptive summary of the sequence lengths used for this benchmarking is illustrated in Table 2. For this analysis we used the Python bindings for the Selenium WebDriver API. In short, our test script, pseudo-randomly selected sequences from the above mentioned fasta file. The number of sequences selected was incremented by one sequence at time, in a loop ranging from 1 to 50. The Selenium WebDriver was used for fasta selections through the PhyloPi web interface the same way a human would have done. Timestamps were taken at each step in the pipeline and this data was used to produce Fig 3 and Table 3. Initial setup of the blastn database for the 11 337 sequences described in table 2, takes approximately 30 seconds.

**Table 2.** Descriptive summary of the sequence lengths from the Los Alamos database used for benchmarking.

| Min | 1<sup>st</sup> Quartile | Median | Mean | 3<sup>rd</sup> Quartile | Max |
| --- | --- | --- | --- | --- | --- |
| 1764 | 3003 | 3012 | 3010 | 3012 | 3176 |

**Table 3.** Time requirements for the various steps in the PhyloPi pipeline.

| Step (Software) | N query sequences | N Total sequences | Time (mm:ss) |
| --- | --- | --- | --- |
| Retrieving related sequences (blastn) | 24 | 110 | 04:24 |
| MSA (MAFFT) | NA | 110 | 21:11 |
| Phylogenetic inference (FastTree) | NA | 110 | 01:12 |
| All other steps combined | NA | 110 | 00:27 |
| Total time | NA | 110 | 27:14 |

**Fig 3.** Regression analysis of benchmarking PhyloPi on the Los Alamos HIV sequences. (a) The number of sequences retrieved by BLAST vs. the number of input (query) sequences. (b) Time (s) vs. the number of input (query) sequences. (c) Time (s) vs. the number of sequences included by BLAST, thus the total number of sequences in the alignment. (d) Time (s) for phylogenetic inference with FastTree vs. the total number of sequences in the alignment.

From all the steps in our pipeline the time requirements for the most time-consuming steps are plotted in Fig 3. All but the alignment step has a linear relationship of time required vs. number of sequences. The multiple sequence alignment has a near quadratic relation (Fig 3c) as can be expected. FastTree2 on the other hand, employs heuristics to change the O(N^2^) problem to a O(N) problem, thus the linearity illustrated in Fig 3d [20].

Many genotyping laboratories are confined to 96 well plates for the sequencing reactions and most use 2 or 3 times coverage of the area of interest. This means that most laboratories will be able to fit 16 or 24 patient samples on a plate considering the target for genotyping is the protease and reverse transcriptase drug resistance relevant area. In our analysis we have included the whole POL coding region which is thus an overestimation. However, including the 5 best blastn retrieved sequences in our analysis we can demonstrate the time requirements for 24 sequences as shown in Table 3.

It is worth mentioning that implementing a combined protease, reverse transcriptase as well as integrase resistance testing assay is possible. This would however require more sequencing primers and reduce the number of patient samples being processed per 96 well plate. Due to the way sequences are retrieved by blastn, our pipeline can handle mixed sequences in the BLAST database. Most laboratories perform a combined protease and reverse transcriptase assay and only do an integrase assay on patients exposed to integrase inhibitors. It is necessary however, to submit queries of different areas separately, i.e. the sequences being submitted for analysis should be capable of producing a sensible alignment. Thus, if a query containing integrase sequences is submitted, blastn will retrieve only sequences of integrase.

## Inter- and intra-patient genetic distances of important coding regions

In order to gauge inter- and intra-patient variability and how to display this in a colour scaled heatmap we again examined sequences from the Los Alamos HIV database. In short, to look at inter- and intra-patient genetic distances of HIV isolates, sequences for the protease and reverse transcriptase coding regions, as a combined sequence (PR_RT), and sequences for the integrase genes were downloaded for the two subtypes, B and C, separately. The fasta files for the 2 subtypes were parsed with python scripts. One script was used to keep only one sequence per patient while another python script was used to put sequences from same patients in to separate fasta files; which we refer to here as patient clusters. This inter- and intra-patient filtering was done based on the Los Alamos HIVdb ‘Patient id’. Any sequence where this data was not available was excluded from the analysis. The inter-patient fasta files were aligned as one file, while the intra-patient clusters were aligned separately by the use of a bash script; again using MAFFT as aligner. A breakdown of the number of sequences and patient clusters is shown in Table 4. As for the phylogenetic inference, for the genetic distance calculations the K80 [24] substitution model was used. The genetic distances from all the clusters were calculated with seqinr and ape as described above.

**Table 4.**
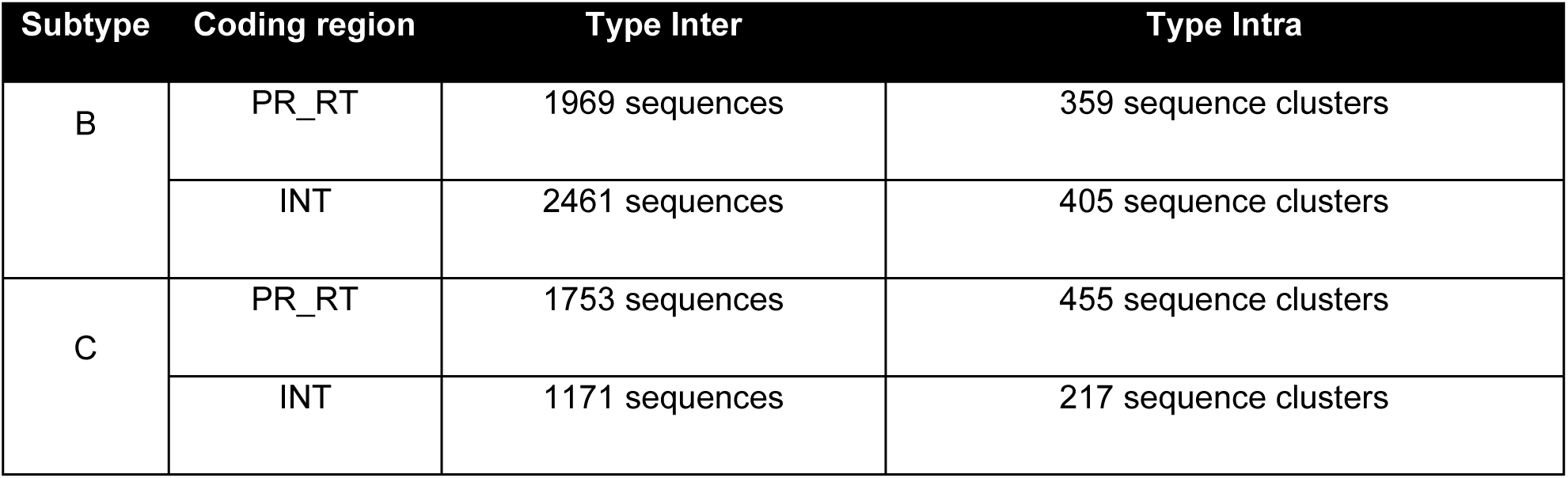
Breakdown of sequences used in the inter- and intra-patient genetic distance analysis.

| Subtype | Coding region | Type Inter | Type Intra |
| --- | --- | --- | --- |
| B | PR_RT | 1969 sequences | 359 sequence clusters |
|  | INT | 2461 sequences | 405 sequence clusters |
| C | PR_RT | 1753 sequences | 455 sequence clusters |
|  | INT | 1171 sequences | 217 sequence clusters |

The violin plots (Fig 4a) shows overlap between types inter- and intra-patient for both subtypes and for both the integrase (INT) as well as the protease reverse transcriptase (PR_RT) coding regions. From the violin plots, it is clear that for type inter-patient the INT coding region is more conserved than the PR_RT coding region.

**Fig 4.** Genetic distance analysis of type inter and intra. (a) Genetic distances depicted in violin graphs of both subtype B and C for patient comparisons, type intra and inter separately showing INT (integrase) and PR_RT (protease reverse transcriptase) coding regions. (b) Logistic regression models showing predicted probability of sequences being of type inter or intra. A probability of one correlates strongly with type intra where a probability of zero strongly correlates with type inter. The dotted lines show the distance intercepts of a 0.5 probability cut-off.

Logistic regression was performed using the R glm library and modeled results are plotted in Fig 4b. From this it is clear that subtype B is more conserved compared to subtype C for the coding regions analyzed here. Also the INT part of the POL gene for both subtypes is more conserved compared to the PR_RT part of the POL gene. Our findings here are in agreement with Li *et al* (2015), who have found that the integrase protein sequence is the most conserved among the 15 proteins they analyzed followed by the rest of the viral enzymes. Comparing the distance intercepts for a predicted probability of 0.5 for either type inter or intra, we can see there is a noticeable difference between subtypes and coding regions. Thus, from the results shown in Fig 4, it is difficult to select one genetic distance cutoff that will fit all subtypes and coding regions. It is also conceivable that the probability of finding transmission clusters will increase as the blastn database increases.

## Discussion

We have demonstrated that this affordable single board computer is capable of performing phylogenetic inference in an impressive short time and is capable of retrieving the most significant sequences from its automatically maintained blastn database without requiring any additional user intervention. Also, the utility of this self-contained pipeline was shown in our case studies. Additionally, PhyloPi provides a search interface for retrieving sequences by name as well as past analysis stored in a SQL database. PhyloPi also provides a tool with which the user can easily map sequences in a fasta file against HIV HXB2 using blastn. This quickly gives a result showing the areas the sequences cover in the POL region, which can be useful to determine whether input sequences are suitable for alignment.

Currently PhyloPi approximates maximum likelihood phylogenetic inference using the K80 (Kimura 80) model [24]. This model assumes equal base frequencies, but different rates of transitions *vs.* transversions. It is very difficult to have an optimal model if the area of the HIV pol gene submitted for phylogenetic inference can change as well as differences in sample batches. Rather than overfitting the data by using too many parameters we use this simple two parameter model. The downside of this, however, is the overestimation of branch lengths. As illustrated in our case studies, PhyloPi were still successful in flagging inspection of sequence results which could then be resolved by further investigation. It is also worth mentioning that all the intermediate files leading to the final result are available for each analysis. The user can intervene at any step. A laboratory might want to validate the use a PhyloPi by using the blastn combined results using their current methods or the already trimmed alignment file for phylogenetic inference using their preferred selection of software and model.

We do not suggest using PhyloPi for finding transmission clusters; rather if two seemingly unrelated patients’ have viral sequences which cluster close together it is possible that the virus was transmitted from one patient to the other, which explains the clustering rather than a contamination issue. This becomes more likely as the blastn database grows.

The logistic regression showed that there is no clear cutoff between inter- and intra-patient HIV sequences and that this is dependent on the subtype and area being sequenced. Furthermore, as the blastn database grows in size, the likelihood of finding closely related sequences from different patients’ increases. There are different sources of sample contamination. Cross contamination between samples may rather create an abnormally high number of mixed bases in the resulting sequence and it is standard practice to repeat these samples. If a sample with a very high viral load contaminated an adjacent sample or a sample in the same batch with a very low viral load, this may not result in large numbers of mixed bases, but these samples will rather have a genetic distance of zero or very close to it. Amplicon carryover contamination could also either result in a high number of mixed bases or identical sequences that are clustered together in time. The true power of PhyloPi is its ability to detect and trigger inspection of sequencing results from patients where there is no apparent trigger for such investigation from routine quality assurance or clinical information provided as illustrated in the section ‘Application of PhyloPi in the HIV drug resistance testing laboratory’, case 1. It is difficult tracking patients in the health sector and especially so in resource limited settings. We typically find patients with various different patient identifiers even when visiting the same health care facility. Despite efforts from our Central Data Warehouse, it is difficult to match patient details such as names, date of birth and address. Patients might use different names depending on language preferences of the patient and health care provider. Furthermore, transcription errors make this even worse. We have found multiple previous sequence results for patients which we otherwise would not have detected. Human error is inevitable and any safety net to catch these mistakes is worthwhile, PhyloPi provides such a safety net.

## Acknowledgments

We thank Dr ED Cason for suggesting the name PhyloPi.

